# Novel R-type Lectin Domain-Containing Cytotoxins Comprise a Family of Virulence-Modifying Proteins in Pathogenic *Leptospira*

**DOI:** 10.1101/2020.05.13.094169

**Authors:** Reetika Chaurasia, Alan Marroquin, Michael A. Matthias, Joseph M. Vinetz

**Affiliations:** Section of Infectious Diseases, Department of Internal Medicine, Yale University School of Medicine, New Haven, Connecticut 06520-8022, USA

**Keywords:** cytotoxin, ricin B-like, lectin, genotoxin, cytopathic effect

## Abstract

Leptospirosis is a globally important neglected zoonotic disease subject to both small scale outbreaks and weather-driven, large-scale epidemics. Due to gaps in our understanding of *Leptospira* biology, pathogenetic mechanisms of leptospirosis remain largely unknown. Previous data suggest that a gene family, PF07598, unique amongst most known bacterial pathogens and encoding so-called “Virulence-Modifying (VM)” proteins, are important virulence determinants. Here, we show that VM proteins are potent cytotoxins, sharing a distinct domain organization while exhibiting varied mechanisms of cellular toxicity. Structural homology searches using Phyre2 suggest that VM proteins are novel R-type lectins containing an N-terminal ricin B chain-like domain. As is known for native ricin B-chain, recombinant full-length **rLA3490** (most highly up-regulated *in vivo*) and an N-terminal fragment, **t3490**, containing a partial ricin B-domain, bound to asialofetuin and directly competed for asialofetuin binding with recombinant ricin B chain. While **t3490** bound to the HeLa cell surface but was neither internalized nor cytotoxic, **rLA3490** bound to the HeLa cell surface, was rapidly internalized, translocated to the nucleus inducing chromosomal fragmentation, and was rapidly cytolethal, providing strong evidence that *Leptospira* VM proteins are *bona fide* cytotoxins. Because monoclonal antibodies impeding cell entry or intracellular trafficking of ricin holotoxin clearly mitigate its toxicity, that VM proteins share binding and intracellular trafficking mechanisms suggests that anti-VM-protein antibody-based (anti-toxin) therapeutics could ameliorate severe complications of leptospirosis thereby improving prognosis. As most VM proteins are restricted to high-virulence *Leptospira* species with some, e.g., LA3490, being exceptionally potent, their level in serum might be a potentially useful indicator of a poor prognosis, thus identifying high risk patients.

**Author Summary:** The PF07598 gene family encoding Virulence-Modifying (VM) proteins in pathogenic *Leptospira* species is associated with severe manifestations of leptospirosis. Structural homology searches indicate that VM proteins contain an N-terminal ricin B chain-like domain, biochemically confirmed in asialofetuin binding and competitive-binding assays suggesting that VM proteins bind to terminal galactosyl residues of this model ricin B domain binding protein. The leptospiral N-terminal ricin B chain-like domain mediated VM protein binding to HeLa cells. Full-length recombinant protein rapidly led to cell death. Amino acid conservation among PF07598 family members at the N-terminal ricin B chain-like domain suggests that VM protein levels in serum might be a useful biomarker for quickly identifying at-risk patients, and that novel “anti-toxin”-based therapeutics could ameliorate severe complications of leptospirosis, both of which remain to be explored.

## Introduction

Leptospirosis is a globally important neglected zoonotic disease subject to both small scale outbreaks and weather-driven, large-scale epidemics, with substantial impact on veterinary and public health. Conservative estimates suggest that the global burden of human disease due to leptospirosis is on par with cholera and typhoid fever (1-3). Annually, more than 1 million cases and 58,900 deaths are estimated to occur globally with case fatality rates ranging from 5-20% (1, 4). Humans become infected after exposure to freshwater or wet soils contaminated by the urine of mammalian reservoir hosts. Clinical presentation varies from an undifferentiated fever to jaundice, renal failure, pulmonary hemorrhage, shock and fulminant death (5-10). Despite informative *in vitro* and small animal models, the molecular, cellular and immunological mechanisms of disease pathogenesis remain unclear (6, 11).

Previously published genomic, pathogenomic and gene expression data suggest that the PF07598 gene family, encoding the so-called Virulence Modifying (VM) proteins, may contribute to the pathogenesis of leptospirosis. VM proteins contain secretory signal peptides (12), and the expression of various PF07598 gene family members is upregulated, both in vitro, under conditions mimicking the in vivo host environment (13), and in vivo in small animal models of acute infection (14). That VM proteins are restricted to group I pathogenic *Leptospira* and expanded in the most highly pathogenic *Leptospira* species and serovars (12, 14), including the cosmopolitan and lethal serovars Copenhageni and Canicola, further suggests that they are involved in pathogenesis.

Based on structural homology searches using Phyre2 (15) that identified, with high confidence, an N-terminal R-type lectin (ricin B-like) domain in the PF07598 gene family, we hypothesized that VM proteins, like ricin, are cytotoxins. First, we tested whether the putative ricin B-like domain in both recombinant full-length, and a truncated protein containing a (partial) ricin B domain of LA3490, the most highly up-regulated PF07598 gene in vivo (14), like ricin B chain, bound to immobilized asialofetuin, a terminal galactosyl-containing glycoprotein, in vitro (16, 17). Next, we determined whether recombinant full-length and truncated recombinant protein LA3490 produced cytopathic effects on cultured HeLa cells. A detailed comparative cross-serovar and cross-species computational analysis was done to assess potential therapeutic potential of anti-VM protein antibodies and biological plausibility of conserved VM protein ricin B domains as vaccine candidates.

## Results

### Domain architecture analysis of VM proteins

Previous work identified a group I pathogen-specific family of paralogous *Leptospira* proteins uninformatively classified as PF07598 (DUF1561) that are expanded (≥12 copies/genome) in highly pathogenic members of the genus, e.g., *L. interrogans* serovar Lai, some of which including serovars Copenhageni and Canicola are cosmopolitan and of particular public health importance. The PF07598 gene family members were not associated with discernable pathogenicity islands, IS elements, nor virulence gene-related operons (12). Previous studies demonstrated that select paralogs were highly upregulated in vivo (14, 18) during mammalian infection and that some were necessary for virulence (14, 19, 20). Serovar Lai encodes 12 paralogs, whereas serovars Copenhageni and Manilae each contain 13 paralogs (Table 1, gene IDs/locus tags and corresponding Uniprot IDs of all PF07598 paralogs found in serovars Lai, Copenhageni and Manilae; those for which knockout mutants are available and have been implicated in virulence are indicated). Both LA3490 (<u>Q8F0K3</u>) and LA0620 (<u>Q8F8D7</u>) are present and highly conserved among all three serovars, with average amino acid identities exceeding 99% (Table 2, and S1 Appendix 1 – 3). Serovars Copenhageni and Manilae, contain an additional ortholog, LIC_10639 (<u>Q72UL8</u>) and LMANV2_170032 (<u>A0A2H1XAY7</u>), respectively, sharing 94% amino acid identity that is absent from serovar Lai.

**Table 1.** PF07598 gene family members in select Group 1 pathogenic *Leptospira*.

| *L. interrogans* serovar Lai strain 56601 |  |  | *L. interrogans* serovar Copenhageni strain 495 |  |  | *L. interrogans* serovar Manilae strain 495 |  |  |
| --- | --- | --- | --- | --- | --- | --- | --- | --- |
| Gene locus | Reference / Protein ID | Amino acid | Gene locus | Reference / Protein ID | Amino acid | Gene locus | Reference / Protein ID | Amino acid |
| LA_3388 | Q8F0V3 | 631 | LIC_10778 | Q72U83 | 631 | LMANV2_260038 | A0A2H1XD57 | 631 |
| LA_0835 | Q8F7V7 | 631 | LIC_12791 | Q72NP1 | 631 | LMANV2_240079 | A0A2H1XD04 | 438 |
| **LA_0591<sup>#</sup>** | Q8F8G6 | 313 | LIC_12985<sup>#</sup> | Q72N53 | 313 | LMANV2_70075<sup>#</sup> | A0A2H1XLT9 | 314 |
| **LA_0589<sup>v</sup>** | Q8F8G8 | 632 | LIC_12986 | Q72N52 | 632 | LMANV2_70078 | A0A2H1XJZ3 | 549 |
| LA_1402 | Q8F6A7 | 641 | LIC_12339 | Q72PX8 | 663 | LMANV2_210058 | A0A2H1XCB0 | 640 |
| LA_1400 | Q8F6A9 | 573 | LIC_12340 | Q72PX7 | 627 | LMANV2_210056 | A0A2H1XC81 | 627 |
| LA_3271 | Q8F166 | 636 | LIC_10870 | Q72TZ4 | 636 | LMANV2_80114 | A0A2H1XKP0 | 638 |
| LA_0934 | Q8F7L0 | 638 | LIC_12715 | Q72NW3 | 638 | LMANV2_320010 | A0A2H1XE97 | 638 |
| LA_0769 | Q8F820 | 602 | LIC_12844 | Q72NJ0 | 639 | LMANV2_240142 | A0A2H1XCY6 | 639 |
| LA_2628 | Q8F2Y3 | 638 | LIC_11358 | Q72SM1 | 638 | LMANV2_150103 | A0A2H1XAR4 | 638 |
| NA | - | - | LIC_10639 | Q72UL8 | 640 | LMANV2_170032 | A0A2H1XAY7 | 638 |
| LA_0620 | Q8F8D7 | 637 | LIC_12963 | Q72N74 | 637 | LMANV2_70050 | A0A2H1XJU7 | 637 |
| LA_3490 | Q8F0K3 | 639 | LIC_10695 | Q72UG2 | 639 | LMANV2_170091 | A0A2H1XB88 | 637 |

NA Not applicable due to a missing ortholog
<sup>#</sup> Signal peptide and complete C-terminal fragment but missing entire N-terminal ricin B chain-like binding domain
<sup>v</sup> Only mutant unable to infect hamster

**Table 2.** Average amino acid identity of LA3490 found in important *L. interrogans* serovars (full-length, N-terminal carbohydrate binding domain and C-terminal putative toxin domain shown).

| Serovars | Lai |  |  |  |  |  |
| --- | --- | --- | --- | --- | --- | --- |
|  | UniProt Protein ID | LA_3490/Q8F0K3 |  | Uniprot Protein ID | LA_0620/Q8F8D7 |  |
| Copenhageni | Q72UG2 | 99.7 |  | Q72N74 | 99.7 |  |
|  |  | 99.6 | 99.7 |  | 99.6 | 99.7 |
| Manilae | A0A2H1XB88 | 97.4 |  | A0A2H1XJU7 | 98.5 |  |
|  |  | 98.7 | 98.4 |  | 97.8 | 98.9 |
| Canicola | A0A1X8WIC7 | 99.7 |  | A0A1R0JW34 | 97.6 |  |
|  |  | 97.8 | 99.7 |  | 97.8 | 97.3 |
| Hardjo subtype *prajitno* | A0A0M4NB61 | 100 |  | A0A0M3TMJ8 | 99.7 |  |
|  |  | 100 | 100 |  | 99.1 | 100 |
| Pomona | A0A163QL98 | 93.5 |  | A0A163P0K6 | 97.4 |  |
|  |  | 97.3 | 97.3 |  | 97.3 | 97.3 |
| Amino acid identity |  | 98.4 |  |  | 98.6 |  |
|  |  | 98.9 | 98.9 |  | 99.1 | 98.9 |

Full length Putative N-terminal ricin B-chain domain Putative C-terminal enzymatic domain

Apart from group I pathogenic *Leptospira*, PF07598 orthologs are found in a number of facultative intracellular bacterial genera including a large family in *Bartonella* (21, 22), and single copies in Campylobacterales (*Campylobacter* spp., *Helicobacter* spp.), *Piscirickettsiaceae*, Actinoplanes and some Vibrionaceae (12). Unlike highly pathogenic group I *Leptospira* (*L. interrogans, L. kirschneri* and *L. noguchii*) and *B. bacilliformis* and *B. australis* that contain 12 or more paralogs per genome, these other genomes encode only single copy homologs with only a few non-*Leptospira* genomes encoding multiple (as many as 4) additional paralogs. Multiple sequence alignment via MAFFT version 7 and protein-disordered structural analysis in Jalview v2.10.5 (S1 Fig) demonstrated that serovars Lai, Copenhageni, Manilae, Canicola, Hardjo and Pomona encode 10 – 12 long, multidomain paralogs (Fig 1A) with domain organization similar to but in reverse orientation of castor bean-produced ricin toxin (23), and distinct from most other bacteria-secreted exotoxins (24-28) that are usually encoded by two or more genes and assembled into multimeric protein complexes (26). All six serovars also contain a shortened PF07598 paralog containing a signal peptide but lacking the N-terminally located ricin B-like binding domain (Fig 1A and below).

**Fig 1.**
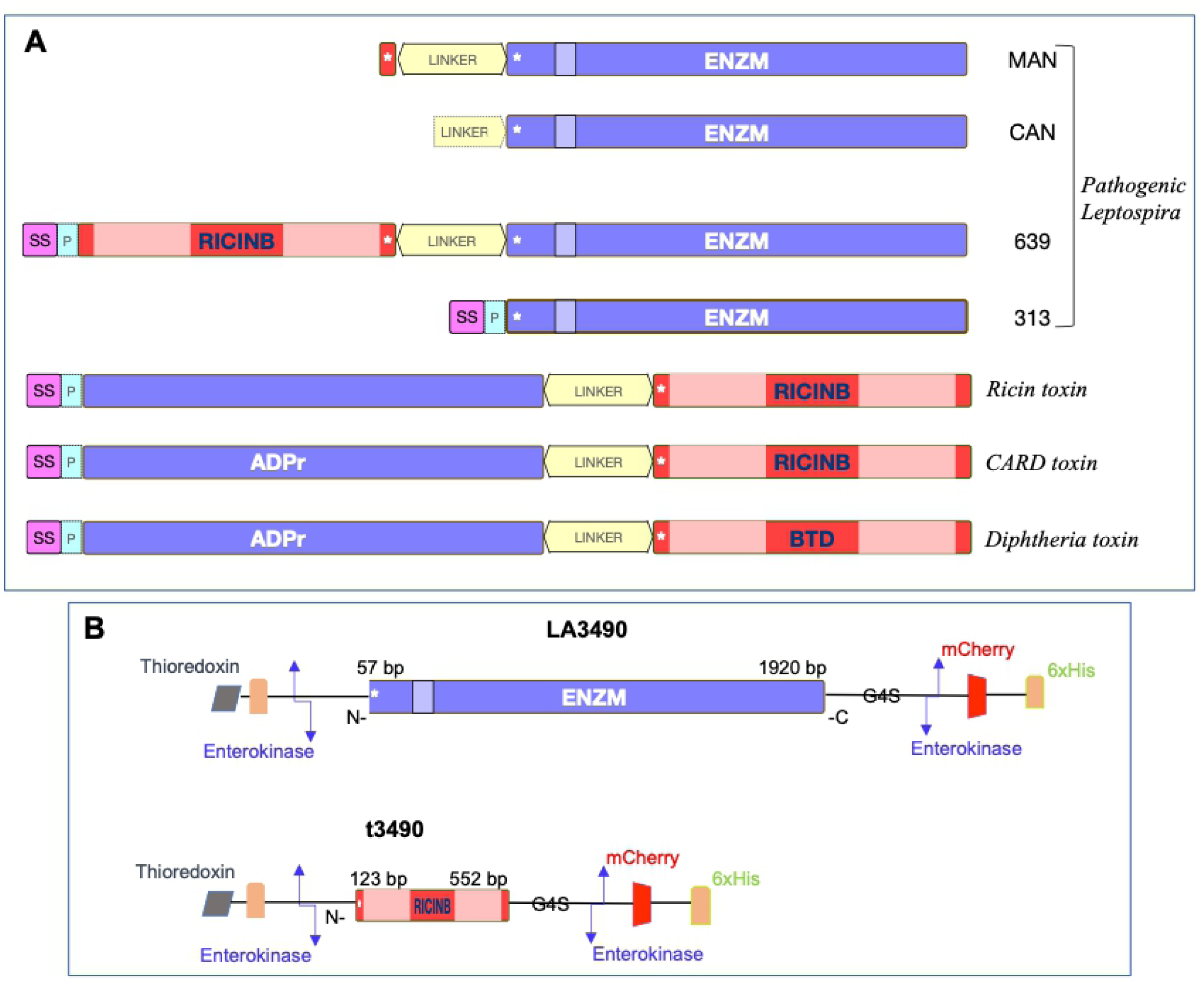
Proposed domain organization of VM proteins, and strategy for cloning of full-length and N-terminal ricin B domain of LA3490. **(A)** Schematic illustration of VM protein domain organization showing the presence of an N-terminal ricin B domain, hydrophobic patch (*) presumed to mediate insertion into endosome membrane prior to the extrusion and release of C-terminal toxin domain to the cytosol. **SS**; signal sequences identified by SignalP. All VM proteins share the LA3490 (<u>Q8F0K3</u>; 639 aa) domain organization depicted, though all serovars also contain a short VM protein variant, e.g., LA0591 (<u>Q8F8G6</u>; 313 aa) and LMANV2_240079 (<u>A0A2H1XD04</u>; 438 aa), containing a SS and complete enzymatic domain (with hydrophobic patch) but lacking the entire N-terminal ricin B binding domain. For reference, the domain organization of plant derived ricin (*Ricinus communis*; <u>P02879</u>, 565 aa), the secreted Community-Acquired Respiratory Distress syndrome (CARDs) toxin of *Mycoplasma pneumoniae* (<u>P75409</u>, 591 aa) and *diphtheria toxin* (*Corynebacterium diphtheriae*; <u>Q5PY51</u>, 535 aa) have been included. Unlike VM proteins, the domain order is reversed in these and other “protoxins” typically encoded by a single polypeptide chain. CARDs and diphtheria toxin belong to the family of ADP-ribosylating toxins; whereas, ricin inactivates the ribosome and inhibits protein synthesis. **(B)** Schematic depicting the organization of the recombinant mCherry fusion proteins used in the current study; **rLA3490** full-length, nucleotide positions 57 - 1920 (minus SS); and **t3490**, 213 - 552, also lacking SS. Recombinant fusions also include a glycine-serine (Gly4S) linker (for flexibility), C-terminal His6 tag (purification), and two internal enterokinase recognition sites.

### Leptospiral ricin B-like domains have carbohydrate binding properties similar to that of ricin B chain

Phyre2 searching (http://www.sbg.bio.ic.ac.uk/phyre2) (15) identified a ricin B-like *β*-trefoil domain in the amino-terminal region of leptospiral VM proteins leading us to hypothesize that, like ricin B, and other ricin B-domain containing bacterial toxins, e.g., *Vibrio cholerae* cytolysin, VCC (29), VM proteins ought to bind to host cell-surface glycoconjugates. We focused here on <u>Q8F0K3</u> (LA3490) because our previous data indicated that it is implicated in virulence and is highly up-regulated in vivo (14). Recombinant <u>Q8F0K3</u> (rLA3490) was expressed in *E. Coli* as an N-terminal fusion with thioredoxin and C-terminal fusion with mCherry and a His_6_ tag to facilitate folding and affinity purification and visualization of the protein using fluorescence microscopy, respectively (Fig 1B). Because native ricin B binds to terminal galactosyl residues of glycoconjugates (30), we used asialofetuin, a terminal galactosyl-containing glycoprotein used for assaying the presence of ricin holotoxin (16, 17, 31), to determine whether rLA3490 would have similar carbohydrate binding specificity. Soluble recombinant proteins, both full-length and truncated, t3490 (i.e. partial ricin B domain, lacking the middle and C-terminal domains (putative translocation and enzymatic)) (Fig 1B), were purified by nickel affinity chromatography, verified by Western immunoblot, and determined to have low levels of endotoxin contamination using a <u>L</u>imulus <u>A</u>mebocyte <u>L</u>ysate (LAL) assay (Fig 2). Binding and ricin B competition assays confirmed that both rLA3490 and t3490 bound to asialofetuin, that binding was mediated by the N-terminal carbohydrate binding domains (CBDs) and that these shared ligand specificity with ricin B (Fig 3).

**Fig 2.**
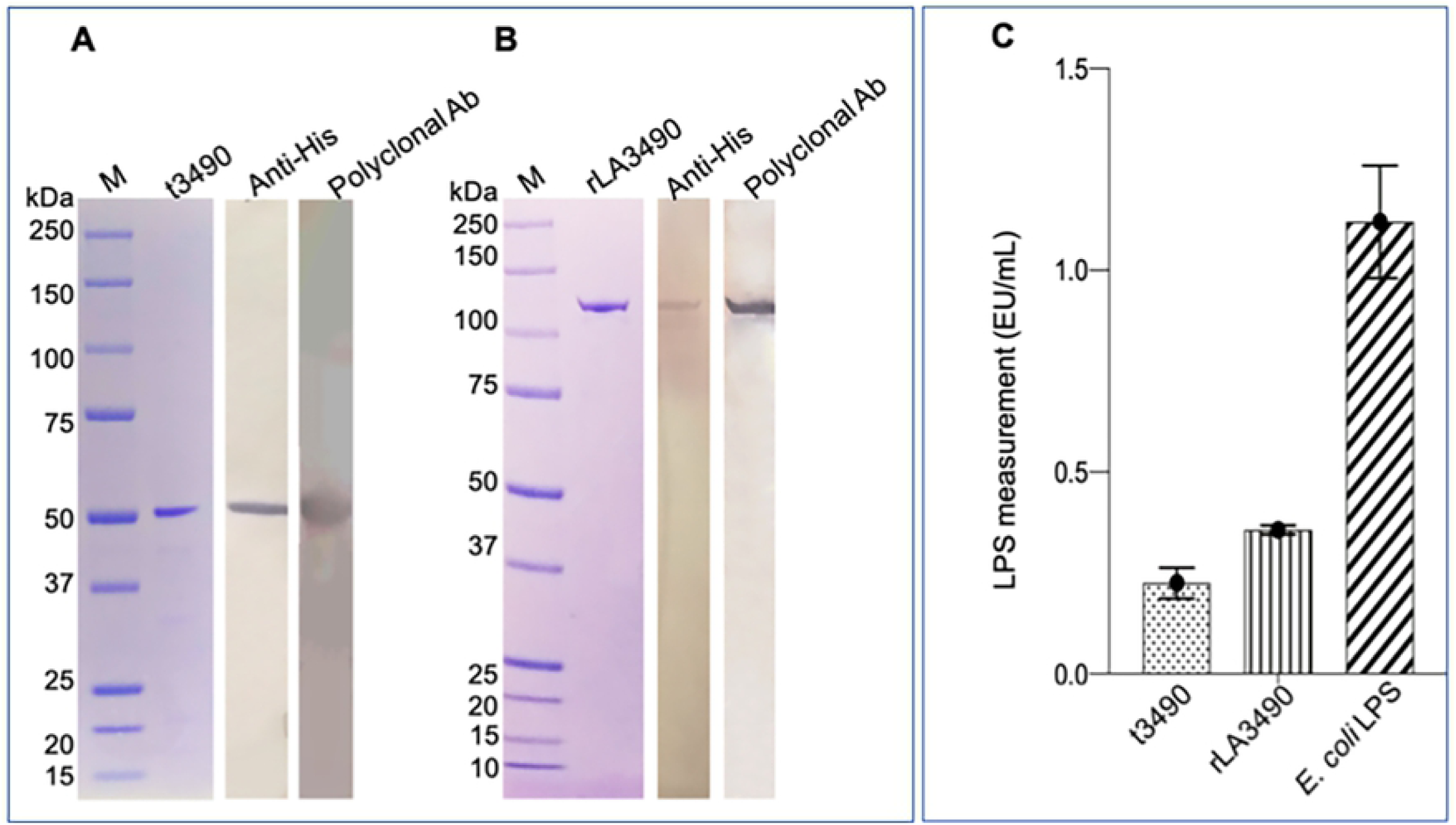
Evaluation of the purity of recombinant t3490 and rLA3490 fusion proteins by Western blot and Limulus Amebocyte Lysate assay. (**A**+**B**) Western blot of purified soluble recombinant proteins confirming the presence single bands of the expected size (**A**, t3490; and **B**, rLA3490). Membranes were probed with anti-His6 and polyclonal anti-LA3490 antibodies (lane two and three, respectively). **M** - molecular weight marker. (**C)** Limulus Amebocyte Lysate (LAL) assay indicating no appreciable endotoxin contamination; *E. coli* LPS used as positive control. Data were visualized in GraphPad Prism v8.

**Fig 3.**
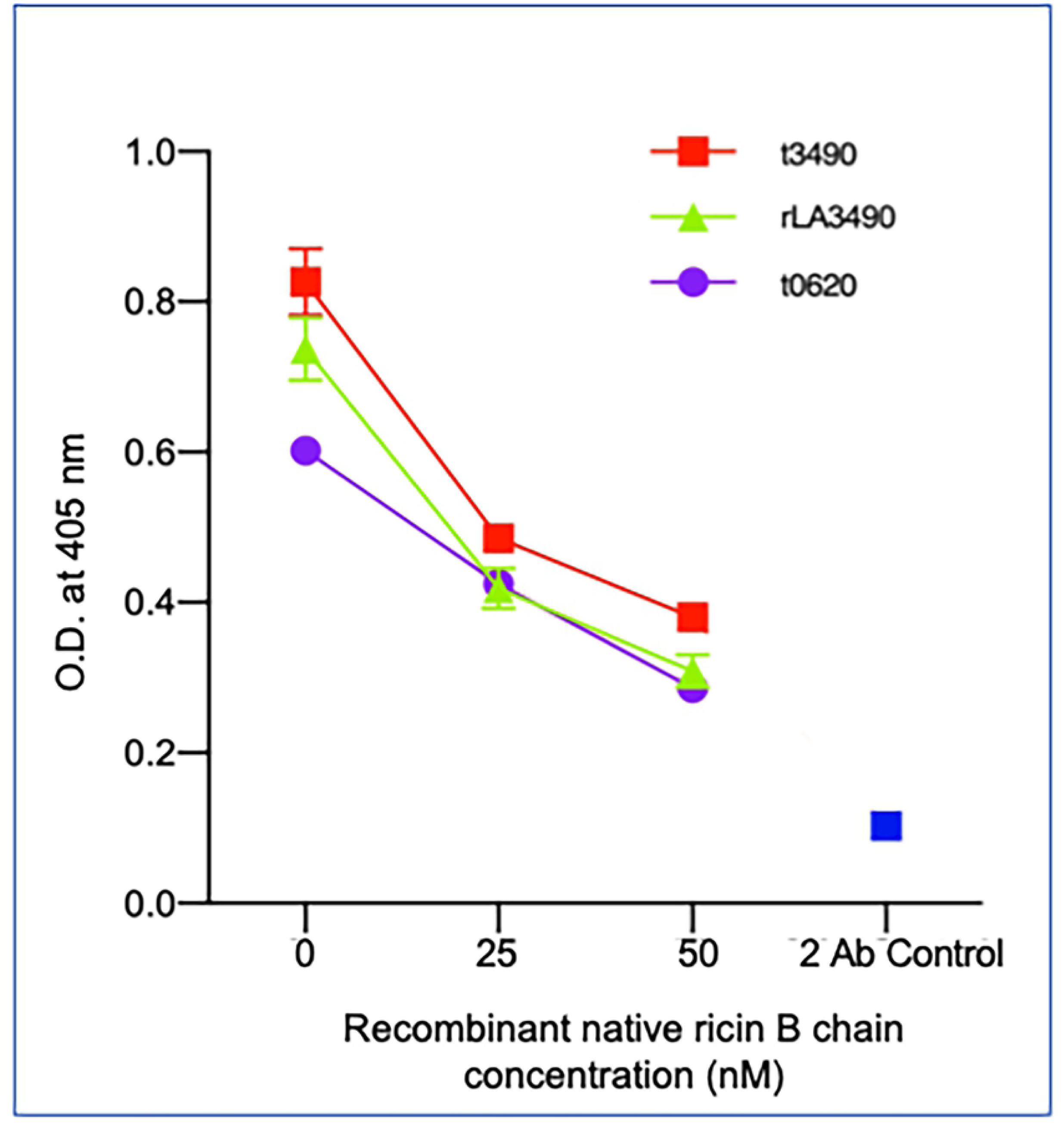
Asialofetuin- and ricin B chain competitive-binding assay. Asialofetuin binding assay confirming that both truncated (**t3490** or **t0620**) and full-length (**rLA3490**) VM proteins bind asialofetuin in the absence (0 nM) of commercially available ricin B chain; and that binding is inhibited by increasing concentrations of ricin B chain (25 nM and 50 nM). Assays were performed in microtiter plates using an ELISA format. Mouse polyclonal anti-LA3490 and anti-LA0620 antibodies (1:1000 dilution) were used as primary and anti-mouse IgG as secondary antibody (used alone as a specificity control). Assays were run in triplicate and experiments repeated at least twice to assess consistency. The mean absorbance (±SEM) were visualized in GraphPad Prism 8 (data from a representative experiment shown).

### Full-length, rLA3490, but not t3490, causes cytopathic effects on HeLa cells

Some bacterial cytotoxins such as those of *Bordetella pertussis, Bacillus anthracis, Pseudomonas aeruginosa, Yersinia pestis, Vibrio cholerae* and *Salmonella typhi*, bind to cell surfaces, become internalized and then exert their effects on one or more intracellular targets (28, 29, 32). Having established that VM proteins contain *bona fide* N-terminal ricin B-like domains, we tested the hypothesis that full-length VM proteins were indeed cytotoxins. Cytopathic effect including cell rounding and blebbing (Fig 4A, and S1 + S2 Video), release of lactate dehydrogenase (Fig 4B), and cell death (Fig 5A) and detachment (Fig 5B), occurred in HeLa cells treated with rLA3490. Such changes were not observed with t3490, bovine serum albumin (BSA) or in untreated HeLa cells. Upon exposure to rLA3490, HeLa cell adhesion was reduced significantly compared to controls (40% vs >80%) (Fig 5B).

**Fig 4.**
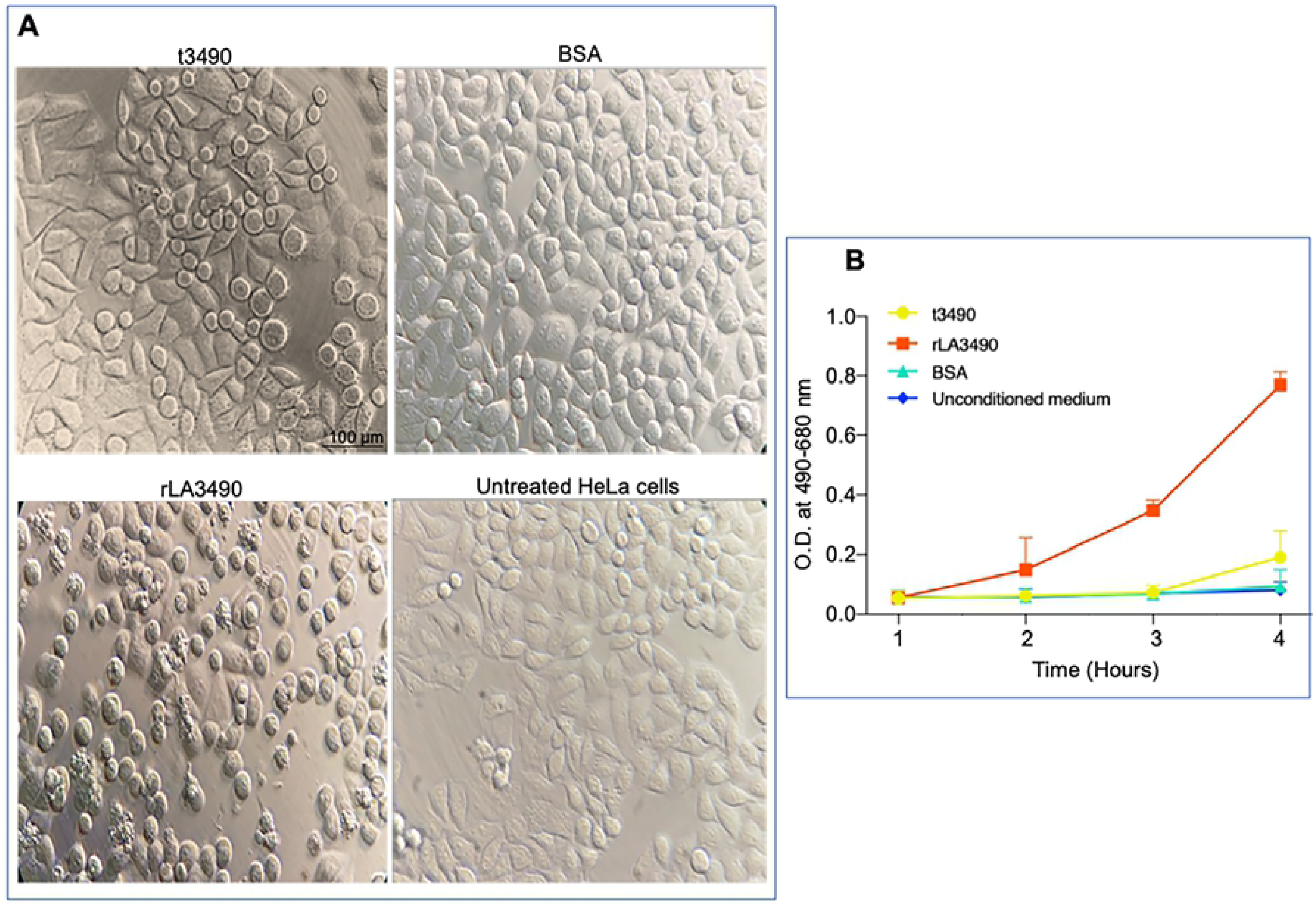
HeLa cell morphology and integrity upon exposure to t3490 and rLA3490. **(A)** Phase contrast images showing alteration of HeLa cell morphology following 4 h exposure to 5 *µ*g/mL of rLA3490, with cell lysis confirmed by the release of lactate dehydrogenase at 1-hour intervals up to 4 h post exposure **(B)**. No such alterations, nor release of LDH, were observed in BSA-treated (5 *µ*g/mL) or untreated cells **(A, B)**. Images were captured at 10x magnification using a Leica DMi8 inverted microscope. Scale bar, 100 *µ*m. Mean absorbance of samples run in triplicate (±SEM) were visualized in GraphPad Prism 8.

**Fig 5.**
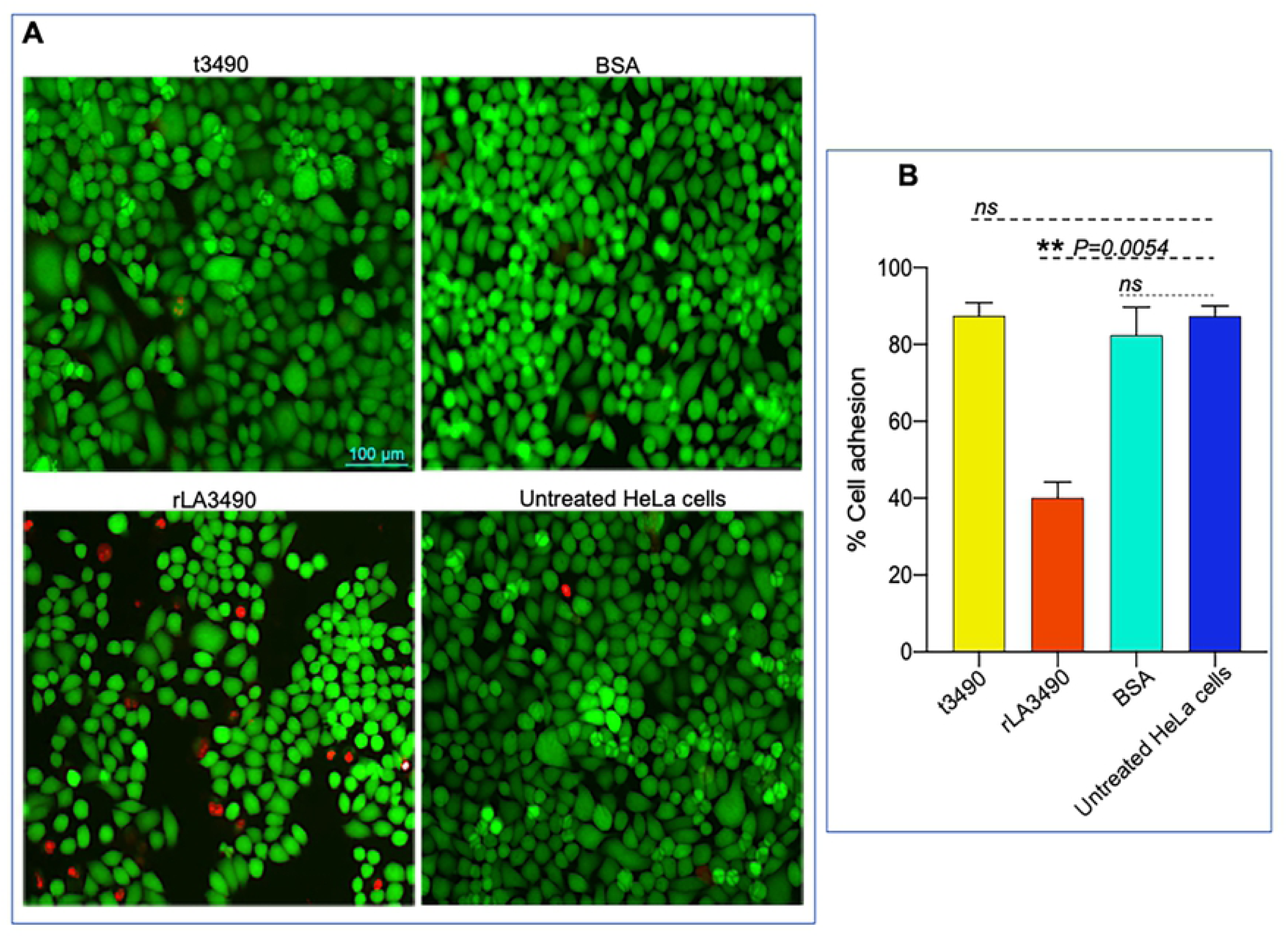
Integrity of HeLa cell monolayers following exposure to t3490 and rLA3490. **(A)** Live/dead staining of HeLa cell monolayers following 4-h exposure to 5 *µ*g/mL t3490 (top left panel) and rLA3490 (bottom left), showing a dramatic decrease of adherent cells and concomitant accumulation of dead cells upon exposure to rLA3490, but not t3490- or BSA-treated or untreated monolayers. Images were captured at 10x magnification using a Leica DMi8 inverted microscope. Scale bar, 100 *µ*m. **(B)** Cell adherence following 4-h exposure to t3490 and rLA3490. Relative to BSA treated and untreated controls, rLA3490, but not t3490, significantly reduced the proportion of adherent cells indicating that N-terminal fragment alone is unable to cause cell death **(A, B)**. The impact of the various treatments was evaluated via Z-test and considered significant when ***p*** < 0.05, **ns** = non-significant.

To further corroborate these findings, live-cell experiments using a transposon mutant of serovar Manilae, M1439, containing a disrupted LA3490 ortholog (LMANV2_1700091, Table 1), which had no cytopathic effect on HeLa cells, and an isogenic wildtype strain that had a pronounced effect (Fig 6). To induce expression of LMANV2_1700091, *Leptospira* cells were grown in liquid EMJH in the presence of 120mM NaCl plus 10% rat serum, conditions previously shown to mimic the host environment (13, 33). Such conditioned *Leptospira* were used to infect HeLa cell monolayers at a <u>M</u>ultiplicity <u>O</u>f Infection (MOI) of 100:1. Whereas wildtype *L. interrogans* serovar Manilae produced cytopathic effect within 2 h, neither M1439 nor the ‘mild’ pathogen, *L. licerasiae* serovar Varillal, which lacks the PF07598 gene family entirely (12), had any noticeable cytopathic effect, even 4 h post-infection (Fig 6), confirming that LA3490 and its orthologs (e.g., LMANV2_1700091) have potent cytotoxic activity.

**Fig 6.**
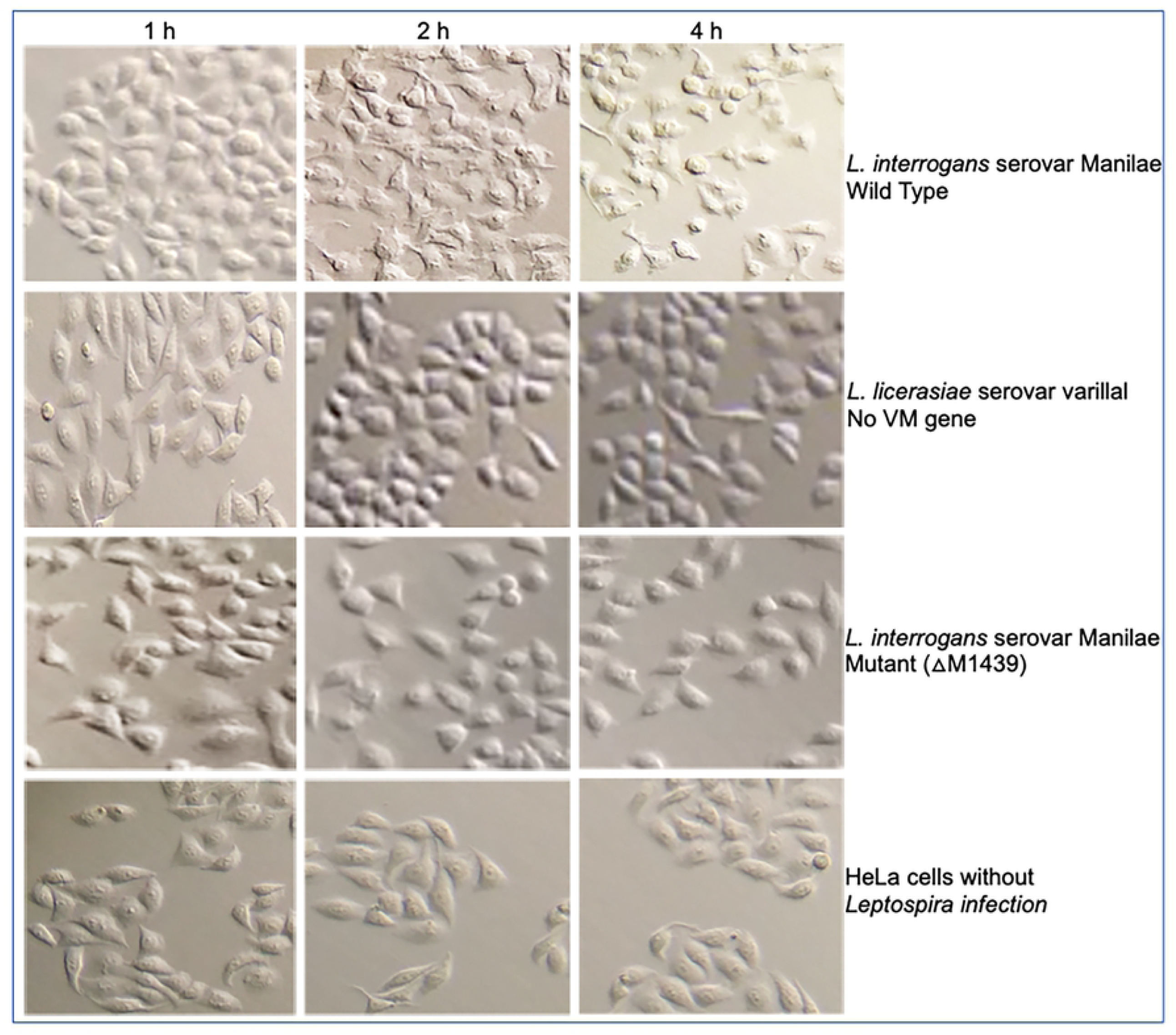
Cytopathic effect of LA3490 knockout mutant strain of *L. interrogans* serovar Manilae and (isogenic) wildtype strain on HeLa cell monolayers. Phase contrast images of HeLa cell monolayers in co-culture with three *Leptospira* strains for up to four hours. Row **<u>1</u>**: transposon mutant, M1439, containing an inactivated LA3490 ortholog, LMANV2_1700091; row **<u>2</u>**: isogenic wild type strain; row 3: low-virulence Group II pathogen, *L. licerasiae* serovar Varillal; and row **<u>4</u>**: uninfected HeLa cell monolayers (control). To induce expression of LMANV2_1700091, *Leptospira* were grown in liquid EMJH supplemented with 120 mM NaCl and 10% rat serum, mimicking the *in vivo* host environment, then used to infect HeLa cell monolayers at a multiplicity of infection of 100:1. Infected monolayers were incubated for up to four hours, with images taken at 1-h intervals for the duration at 10x magnification using a Leica DMi8 confocal microscope. Scale bar, 100 *µ*m.

### Full-length VM protein LA3490 is internalized by HeLa cells and is translocated to the nucleus

To test whether rLA3490 (and VM proteins in general) are internalized by HeLa cells, rLA3490- and t3490-mCherry fusion proteins were visualized by super-resolution confocal microscopy (Leica SP8 Gated STED 3X). Like t3490, rLA3490 bound to the cell surface, but the full length protein was internalized and translocated to the nucleus. Maximum binding occurred 30 – 60 minutes post-exposure, with internalization, translocation and nuclear degradation evident from 30 min onwards; t3490 bound to the surface of HeLa cells but was not internalized (Fig 7).

**Fig 7.**
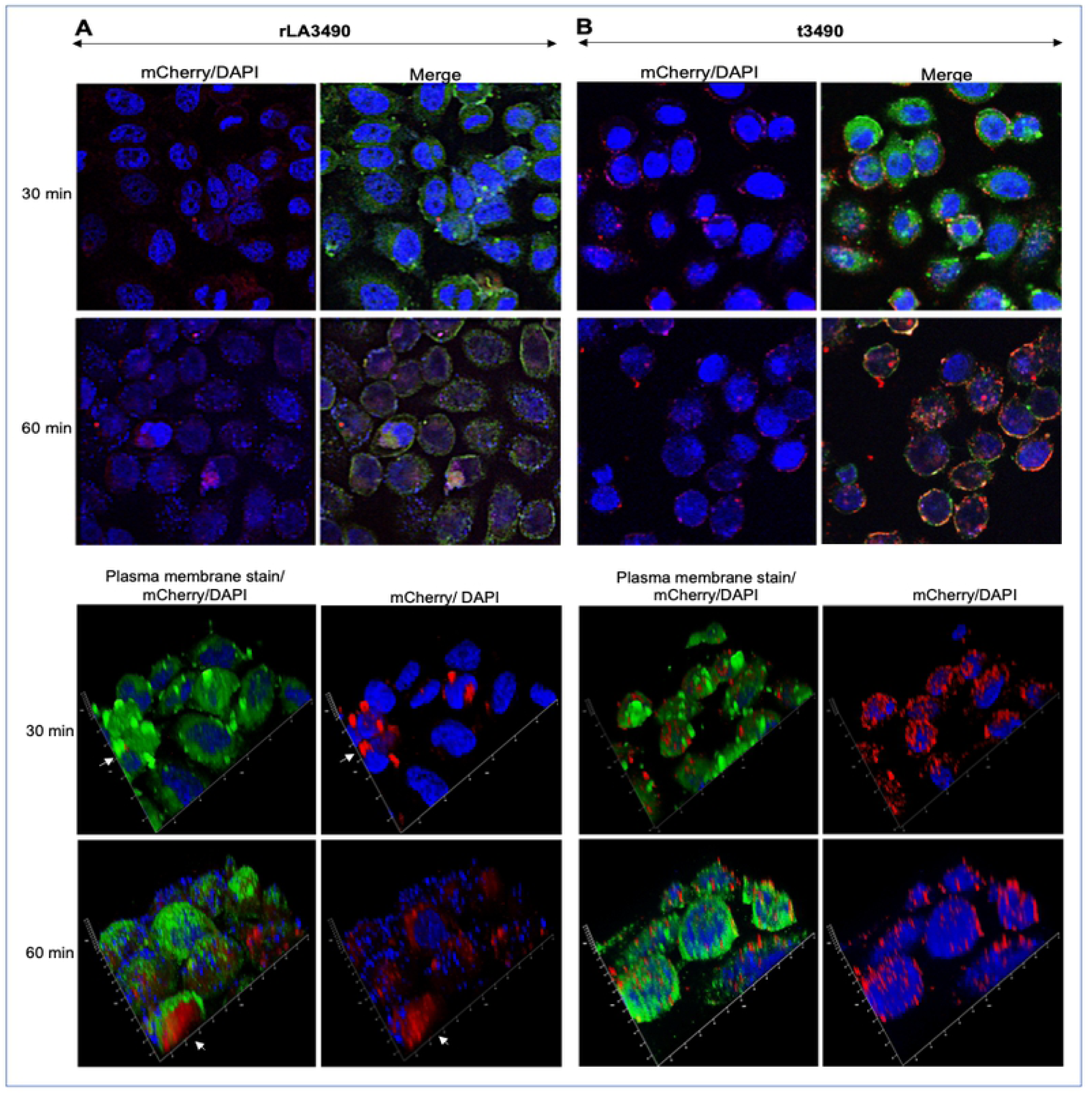
Intracellular trafficking and localization of rLA3490 in HeLa cells. Confocal images show that whereas both **mCherry-rLA3490** and -**t3490** fusion proteins bind to the cell surface (**A** and **B**, respectively), only rLA3490 is internalized. Z-stack images showing **mCherry-rLA3490** fusion was internalized from 30 min onwards, with nuclear translocation and chromosomal degradation (evinced by patchy DAPI staining; **A**, panel four, lower right) evident within 60 minutes. Monolayers were exposed to 5 *µ*g/mL recombinant fusion protein or BSA for 30 minutes and 60 minutes (or were left untreated). Cells were washed, stained with CellMaskTM Green Plasma Membrane Stain, and then mounted with ProLong™ Gold Antifade Mountant +DAPI. Images were captured at 100x using appropriate filters (blue, DAPI; green, plasma membrane; and red, mCherry fusions).

### Full-length VM protein rLA3490 induces actin depolymerization in HeLa cells

Internalization of rLA3490 induced depolymerization of actin filaments producing cell rounding from 1 h onwards after treatment (Fig 8A). The morphology of HeLa cells treated with t3490 (Fig 8B) or BSA (Fig 8C), and untreated cells (Fig 8D) remained unaltered. The mechanism(s) by which leptospiral VM proteins, such as LA3490, perturb actin polymerization merits further investigation.

**Fig 8.**
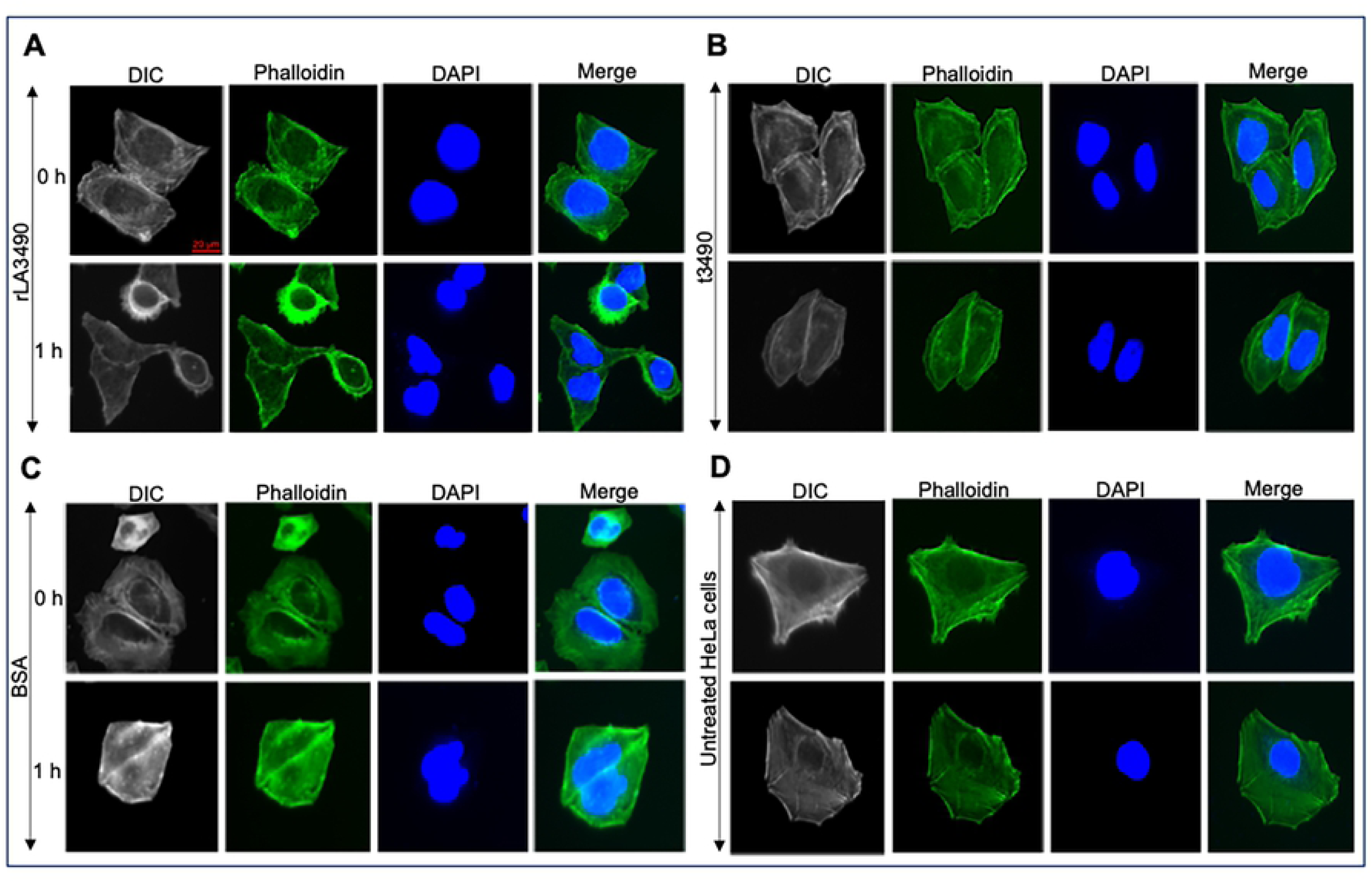
Full-length LA3490 mediated depolymerization of F-actin of HeLa cells. HeLa cell monolayers were incubated with 5 µg/mL of each of rLA3490 **(A)**, t3490 **(B)**, and BSA **(C)** up to 1 h. Monolayers were fixed with 4% paraformaldehyde for 30 minutes and then followed by washes, 0.1% Triton X-100 in PBS was added to each well for 5 minutes. Monolayer was incubated with phalloidin Alexa_488nm conjugate at room temperature for 30 minutes in dark, washed twice and then mounted with ProLong™ Gold Antifade Mountant with DAPI for 10 minutes. Images were captured using a Leica DMi8 confocal microscope with appropriate filters (Alexa_488nm (green), DAPI (blue)) at 40x magnification. Untreated HeLa cells served as control **(D)**. Scale bars 20 µm.

### Distribution of VM paralogs among medically important Leptospira serovars, as potential prognostic biomarkers and implications for biotherapeutics

Based on publicly accessible sequence data, it is clear that VM proteins have an extremely limited taxonomic distribution. Among the organisms known to possess VM proteins, three species of *Leptospira* (*interrogans, kirschneri* and *noguchii*), which comprise the most highly pathogenic *Leptospira*, and two species of *Bartonella* (*bacilliformis* and *australis*) are exceptional in that their genomes contain 12 or more distinct VM paralogs, whereas most other species including other, less virulent, group I pathogenic *Leptospira* encode at most 5 paralogs. Previous comparative whole genome analysis of all (at the time) recognized pathogenic *Leptospira* species hinted at a complex series of gene duplications/deletions underpinning the uneven the distribution of VM proteins amongst *Leptospira*. To further explore this phenomenon with an eye towards biomarker discovery and the development of novel therapeutics, we performed a more focused analysis incorporating the improved understanding of VM protein domain organization (targeting vs enzymatic) uncovered here. Manually curated multiple sequence alignments (S1 Fig) were used to produce detailed pairwise distance matrices of full-length proteins (S1 Appendix 1), and isolated lectin (S1 Appendix 2) and toxin domains (S1 Appendix 3) from six public-health important *L. interrogans* serovars Lai (field mouse reservoir, and reference genome), Copenhageni (rat-borne), Canicola (dogs), Hardjo (cattle), Manilae (rat-borne) and Pomona (broad host range including pigs) that allow us to draw important conclusions. First, as might be expected, the N-terminally located ‘targeting’ lectin domains are on average more conserved (78% amino acid identity, amino acid identity) than the C-terminal toxin domains (63% amino acid identity), consistent with our belief that VM proteins are essentially lectin cytotoxins. Second, the range of targeting and toxin functionalities varies amongst serovars, with both serovar-related duplications of certain lectin domains and lectin domain-toxin domain combinations. For example, both the N- and C-terminal domains of LA3490 are present and highly conserved (98.7% and 98.6% amino acid identity, respectively) in all six serovars analyzed: Lai, Copenhageni, Canicola, Hardjo, Manilae and Pomona. Whereas the full-length Lai and Copenhageni VM proteins are highly conserved between each other in both lectin (99.6% amino acid identity) and toxin (99.7%) domains, in serovar Canicola, for example, the LA3490 toxin domain (97.6% amino acid identity) is linked to a different lectin domain shared with Q72UL8 (LIC_RS03300) (97.8% amino acid identity), which itself occurrs in tandem with a distinct toxin domain (61.6% amino acid identity). In some paralogs, e.g., Q72U83 and Q72NP1, the lectin domains are 100% conserved, but linked to distinct toxin domains (70.7% amino acid identity), suggesting that apart from amino acid sequence variation in either domain, domain shuffling to produce varied lectin and toxin domain combinations could also contribute to the diversity of *Leptospira* VM proteins.

Based on the dramatically differing response of rLA3490- and rLA0620-treated HeLa cell monolayers, the observed sequence variation in the toxin domains could impact potency and/or mechanisms of toxicity. It is also evident that some serovars, e.g., Lai and Hardjo, and Canicola and Pomona have nearly perfect (>99% amino acid identity) conservation of their full-complement of lectin domains and may have similar cellular targets.

## Discussion

Here we demonstrate that *Leptospira* Virulence Modifying (VM) proteins (12, 14) are *bona fide* R-type lectin domain-containing cytotoxins, virtually unique to pathogenic *Leptospira*, some of which rapidly kill mammalian cells after first binding to the cell surface, internalization and nuclear translocation. VM proteins produce cytopathic effect including cell rounding, cell membrane blebbing, pedestal formation, actin depolymerization, nuclear fragmentation and ultimately cell lysis and death. Most contain an N-terminal fragment containing an empirically verified carbohydrate binding domain (CBD) sharing binding specificity with ricin B chain for terminal galactosyl residues of glycoconjugates that likely mediates cell binding/targeting and internalization, and a more variable C-terminal domain (CTD) responsible for intracellular trafficking and cytotoxicity. The high amino acid identity of the N-terminally located ricin B-like domain enables grouping of *L. interrogans* serovars into host cell tropic or virulence groups based on pairwise comparisons of the full complement of N-terminal CBDs or C-terminal CTDs, respectively, based on their respective VM paralogs. The finding that certain VM protein homologs induce cytopathic effect is the first definitive evidence that certain serovars are intrinsically more virulent than others and therefore of heightened clinical and public health significance.

Like ricin, for which toxicity and pathology are clearly linked and route dependent (inhalation leading to severe respiratory compromise being most lethal), site-specific expression of select VM family members could be a parsimonious explanation of the famously poorly understood ‘protean’ manifestations of severe leptospirosis. Indeed, efforts to understand the molecular and cellular pathogenesis of leptospirosis remains in its infancy, and approaches to prevent leptospirosis or ameliorate its pathogenesis are predicated on mechanistic understandings of the biology of *Leptospira*-host interactions. For example, pulmonary hemorrhage and refractory shock are particularly important clinical manifestation of leptospirosis (34-41). Indirect evidence—that these serious manifestations are ameliorated by hemodialysis/hemofiltration (42, 43)—suggests that there may be circulating soluble toxin(s) in leptospirosis. Histopathological analysis of lung tissues in severe pulmonary leptospirosis syndrome do not find intact *Leptospira* (44), but rather damage to alveolar epithelial and activation of endothelial cells, which might explain deposition of immunoglobulin and complement as secondary events (45-47). The data we present here build on. our previously published observations (12-14) that showed *Leptospira* VM proteins to be major virulence factors, potentially involved in the molecular and cellular pathogenesis of leptospirosis. And crucially, new monoclonal antibody based ‘biotherapeutics’ targeting these seemingly potent *Leptospira* R-type cytotoxins could revolutionize patient care and could ameliorate severe manifestations of leptospirosis.

Extensive studies using transposon mutants indicate have shown that multiple distinct VM proteins, including LA0589, contribute to lethal disease in an experimental hamster model (20, 37). The present work is consistent with previous findings that LA3490 is among the most highly upregulated PF07598 family members *in vivo* (14). Phyre2-based *in silico* analysis indicated that VM proteins contain ricin B domains (R-type lectins), which was confirmed experimentally using recombinant full-length VM protein (rLA3490) and N-terminal fragments containing isolated ricin B domains (t3490, t0620). R-type lectins belong to a superfamily of proteins containing a CBD named for and structurally similar to ricin B chain and are found in plants, animals, and bacteria (30). Ricin and its B chain (and other R-type lectins) bind to terminal galactoses or other related glycans of a diverse range of host cell surface glycoconjugates, which facilitates translocation and internalization of the ricin A chain into target cells, resulting in cell death via inhibition of protein synthesis (48-51). Likewise, most potent bacterial toxins such as Shiga, diphtheria and pertussis toxins mediate cell death either by ADP-ribosylation of 28S rRNA or by inactivation of elongation factor 2 (52-55). However, the less well-studied genotoxins, e.g., cytolethal distending toxins (CDTs), are endonucleases that exert their toxic activity following nuclear translocation, similar to LA3490.

While we demonstrated that ricin B domains of distinct VM proteins (LA3490 and LA0620) bound to immobilized asialofetuin (16, 56, 57), native target ligands and cellular targets are yet to be defined. Second, differences in cytopathic potential among VM proteins and molecular pathways by which these proteins exert their biological effects remain to be explored, though these initial experiments clearly indicate that toxicity is mediated by the CTD and is likely to differ among various VM family members because of substantial amino acid sequence variation in the C-terminal half of the molecule. Third, while sequence variation and domain shuffling account for the VM protein diversity, we do not yet understand the reasons for the expansion of PF07598 paralogs in *L. interrogans, L. kirschneri* and *L. noguchii* (14) compared to other pathogenic *Leptospira* species (12, 14), the uneven distribution of VM paralogs, which are clearly non-redundant (12), among medically-important serovars would imply that necessary targeting and toxin functionality are serovar-dependent with host adaptation a potential eco-evolutionary driver.

## Materials and Methods

### Bacterial strains

*Leptospira* were maintained at 30°C in semi-solid **<u>E</u>**llinghausen, **<u>M</u>**cCullough, **<u>J</u>**ohnson and **<u>H</u>**arris medium (EMJH, BD Biosciences, USA) (58). Isogenic, wild type *L. interrogans* serovar Manilae strain L459 (National Veterinary Services Laboratory, Ames, IA, USA), mutant M1439 (transposon mutant LMANV2_170091) (20) and *L. licerasiae* VAR010 (isolated from mild leptospirosis case in Peru) were cultured in liquid EMJH medium to a cell density of ∼2 × 10^8^ cells/mL supplemented with 120 mM NaCl and 10% rat serum (Rockland Immunochemicals, USA) to simulate the in vivo host environment and to induce virulence gene expression (59). *Leptospira* cells were harvested by centrifugation at 18,514 *g* for 20 min at 4°C (Eppendorf, USA), washed twice with cold 1X PBS, pH 7.4 (AmericanBio, USA), and then counted by dark-field microscopy via a Petroff-Hausser counting chamber (Fisher Scientific, USA).

### Mammalian cell culture

HeLa cells were obtained from the **<u>A</u>**merican **<u>T</u>**ype **<u>C</u>**ulture **<u>C</u>**ollection (ATCC, USA) and maintained in **<u>D</u>**ulbecco’s **<u>M</u>**odified **<u>E</u>**agle **<u>M</u>**edium (DMEM; Sigma-Aldrich, USA), supplemented with 10% fetal bovine serum and 1% antibiotic-antimycotic solution (penicillin, 100 units/mL; streptomycin, 100 *µ*g/mL and amphotericin, 25 *µ*g/mL; Invitrogen, USA) at 37°C in a humidified atmosphere containing 5% CO_2_. Antibiotic-containing medium was replaced with fresh, antibiotic-free medium prior to experimental infection with *Leptospira*, which were done at a MOI of 100:1 for 4 h (60).

### In silico analysis

To improve functional classification of PF07598, nucleotide sequences of LA3490 and LA0620 were uploaded to the online Phyre2 server (http://www.sbg.bio.ic.ac.uk/phyre2) (15), which utilizes structural information to detect remote homologs. In addition, coding sequences of all full-length (excluding pseudogenes) PF07598 gene family members in *L. interrogans* serovars Lai strain 56601^T^ [12 paralogs], Cophenhageni L1-130 [13], Canicola [13], Hardjo subtype prajitno Norma [12], Manilae strain L495 [13], Pomona [12], and *L. kirschneri* Pomona [10] were downloaded from Uniprot (http://www.uniprot.com) and aligned using the MAFFT v7 ***einsi*** algorithm with default parameters (http://mafft.cbrc.jp/alignment/software). Poorly aligned regions were refined manually in Jalview v2.10.5, which was also used to predict secondary structure (JPred with default parameters (http://www.compbio.dundee.ac.uk/jpred/jalviewWS/service/JPred)), and to identify globular domains (GlobPlot with default parameters (http://globplot.embl.de)).

### Plasmid constructs and cloning

The gene sequences of LA3490 (Gene Bank Accession number; <u>NP_713670.1</u>, 1920 bp) and LA0620 (<u>NP_710801.1</u>, 1914 bp) were retrieved from NCBI (https://www.ncbi.nlm.nih.gov). Sequences of full-length LA3490 without signal peptide (57 bp – 1920 bp) and N-terminal ricin domain of LA3490 (123 bp – 552 bp) were codon-optimized and fused with full-length mCherry (<u>AST15061.1</u>) as a fluorescent tag (708 bp) along with a glycine-serine hinge (Gly4S) and flanking N- and C-terminally located enterokinase recognition sites (Fig 1B). Synthetic genes were cloned into pET32b (+) (Gene Universal Inc., USA), and verified by sequencing.

### Recombinant protein expression and purification

Because PF07598 gene family members are cysteine-rich, recombinant proteins were expressed in SHuffle**®**T7 competent *E. coli* cells (New England Biolabs, USA), due to their capacity to promote disulfide bonds in the cytoplasm ensuring proper protein folding. Transformants were sub-cultured into Luria-Bertani (LB) medium containing 100 *µ*g/mL ampicillin. When cultures had reached an OD of 0.6, expression was induced at 16°C and 250 rpm for 24 h via addition of 1 mM isopropylthio-*β*-D-galactoside (IPTG; Sigma-Aldrich, USA). Following induction, cells were pelleted then lysed in CelLytic™ B (Cell Lysis Reagent; Sigma-Aldrich, USA) containing 50 units benzonase (Sigma-Aldrich, USA), 0.2 *µ*g/mL lysozyme, non-EDTA protease inhibitor (Roche, USA) and 100 mM PMSF (Sigma-Aldrich, USA) for 1 h at 37°C. Lysates were then centrifuged at 4°C and 18,514 *g* for 10 min, supernatants and pellets were separated, and protein concentration determined by BCA (Pierce™ BCA Protein Assay Kit, Thermo Scientific, USA). Proteins were analyzed by 4-12% bis-tris sodium dodecyl sulfate-polyacrylamide gel electrophoresis (SDS-PAGE). Purification was done using 5 mL pre-packed Ni-sepharose AKTA Hi-TRAP column (GE Healthcare, USA) equilibrated (100 mM NaH_2_PO_4_, 10 mM Tris-HCl, 25 mM imidazole, pH 8.0). Bound protein was eluted in buffer containing 500 mM imidazole, pH 8.0. Pooled eluates were concentrated using a 30 kDa amicon ultra-centrifugation tube (Merck Millipore, Germany) and washed two to three times with 1X PBS pH 7.4 at 4°C. Purified soluble recombinant proteins were tested for bacterial endotoxins using the **<u>L</u>**imulus **<u>A</u>**mebocyte **<u>L</u>**ysate (LAL) assay kit (Invitrogen, USA).

### SDS-PAGE and Western immunoblot analysis

SDS-PAGE was performed according to the method of Laemmli (61). Proteins were stained with Coomassie Brilliant Blue R-250. For Western immunoblot analysis, proteins were transferred onto a nitrocellulose membrane. The membrane was blocked with 5% nonfat dry milk in 1X TBST (TBS + 1% Tween 20) buffer (AmericanBio, USA) for 2 h, and then probed with either mouse anti-His monoclonal antibody (1:2,000 dilution; Santa Cruz Biotechnology, USA) or mouse anti-LA3490 polyclonal antibodies and mouse anti-LA0620 polyclonal antibodies (1:1,000 dilution), respectively. After washing with TBST, the blot was incubated for 2½ h with alkaline phosphatase-conjugated goat anti-mouse IgG (H+L) as the secondary antibody at dilution of 1:5000 dilution (KPL, USA). The blot was developed with ready-to-use 5-bromo-4-chloro-3-indolyl phosphate and nitroblue tetrazolium solution (BCIP/NBT; KPL, USA).

### Asialofetuin binding and ricin B chain competitive binding assay

Binding assays were done using Immulon® 2HB flat-bottom microtiter plates (Thermo Fisher Scientific, USA). Plates were coated with asialofetuin (250 ng/100 *µ*L in carbonate-bicarbonate buffer, pH 9.4), incubated at 4°C for overnight, and then blocked with 5% non-fat skimmed milk in 1X TBST for 2 h at 37°C. After blocking, rLA3490 and t3490 were added separately at a concentration of 50 nM (in 1X TBST). Plates were incubated for 2 h; and washed three times with TBST prior to incubation for 1 h with anti-LA3490 polyclonal antibodies (1:1000 in TBST). Bound rLA3490/t3490 was quantified as follows: plates were incubated with goat anti-mouse IgG (1:5000; KPL, USA) for 1 h, washed thrice with TBST and developed with p-Nitrophenyl phosphate (1-Step™ PNPP Substrate Solution; KPL, USA). The reaction was stopped with 2 M NaOH, and absorbance was read at 405nm on a SpectraMax® M2e Microplate Reader with preinstalled SoftMax®Pro 5.2 (Molecular Devices, USA).

For competitive binding assays, plates were pre-incubated with either 25 nM or 50 nM recombinant ricin B chain (Vector Laboratories, USA) for 2 h before addition of 50 nM of rLA3490 or t3490 and a final 2 h incubation. Bound rLA3490/t3490 was quantified as described in the preceding paragraph.

### rLA3490-mediated HeLa cell cytotoxicity

HeLa cells (35,000 cells/200 *µ*L) were seeded in 8 well chamber slides (LabTek, USA) and incubated at 37°C in a humidified atmosphere containing 5% CO_2_ for 24 h. Cells were treated with 5 *µ*g/mL of rLA3490 or t3490 for up to 4 h; BSA (5 *µ*g/mL) and untreated HeLa cells served as controls. Images were captured at 10X magnification using a Leica DMi8 inverted microscope (Leica Microsystems, Germany). Adherent cells, before and after 4 h exposure to either rLA3490 or t3490, were counted using Leica Application Suite X (Leica Microsystems, Germany).

### Live/dead and F-actin staining, and LDH assay

HeLa cells were treated with 5 *µ*g/mL of rLA3490 or t3490 for 4 h. The monolayer was washed twice with 1X PBS, pH 7.4. Two hundred microliters of 2 *μ*M calcein AM/4 *μ*M ethidium homodimer-1 in PBS (Live/Dead® Viability Kit, Invitrogen, USA) were added to the wells and plates incubated for 30 min in the dark. Monolayers were washed with PBS pH 7.4 to mitigate non-specific, background fluorescence. BSA and untreated HeLa cells were used as controls. Images were taken using a Leica DMi8 microscope at 10x magnification with appropriate excitation and emission filters for green (live cell) and for red (dead) fluorescence. In addition, cell damage was quantified by assaying the release of lactate dehydrogenase (CyQUANT™ LDH Cytotoxicity Assay, Invitrogen, USA).

For F-actin staining, cell monolayers were exposed for up to 1 h, washed twice with PBS, pH 7.4, and then fixed with 4% paraformaldehyde (Sigma-Aldrich, USA) for 30 minutes at room temperature. Following aspiration of the fixative, monolayers were washed twice with PBS, then 0.1% Triton X-100 in PBS was added to each well for 5 minutes prior to repeat washes with PBS. Monolayers were incubated with phalloidin Alexa_488nm conjugate (Invitrogen, USA) at room temperature for 30 minutes in the dark per manufacturer’s directions. Nuclei were stained with 0.1 *µ*g/mL of ProLong™ Gold Antifade Mountant with DAPI (Invitrogen, USA) for 10 minutes. All images were taken using a Leica DMi8 microscope with appropriate filters (Alexa_488nm (green), DAPI (blue)) at 40x magnification.

### Internalization of rLA3490 by HeLa cells

HeLa cells were seeded in 8 well chamber slides (LabTek, USA) and incubated as described above. Monolayers were treated with 5 *µ*g/mL of rLA3490 or t3490 for up to 60 minutes, washed twice with PBS, and then stained with CellMask™ Green Plasma Membrane Stain (Invitrogen, USA) per manufacturer’s directions. Images were taken using a Leica SP8 Gated STED 3X super resolution confocal microscope (Leica Microsystems, Germany) at 100x with appropriate filters.

### Statistical analysis

All experiments were performed in triplicate. Results were expressed as mean and standard deviation and non-parametric *t*-test was used to assess statistical significance in Graph Prism 8.

## Acknowledgments

The work reported benefitted from the use of a (knockout) mutant strain of *L. interrog*ans serovar Manilae containing a transposon inactivated LA3490 ortholog that was a kind gift of Ben Adler, Gerald Murray and Mathieu Picardeau.

## Supporting Information

**S1 Figure**. Amino acid multiple sequence alignment of all full length PF07598 paralogs present in serovars Lai, Copenhageni, Canicola, Hardjo and Pomona.

**S1 Appendices (1-3)**. Pairwise distance matrices of full-length (S1 Appendix 1), N-terminal ricin B chain-like carbohydrate binding domain (S1 Appendix 2) and C-terminal toxin domain (S1 Appendix 3). Pairwise distances of Lai, Copenhageni and Canicola have been conditionally formatted, and all others de-emphasized for easier viewing. Cells colored shades of green indicate low amino acid identity —darker shades = lowest; whereas those colored shades of blue indicate higher amino acid identity —darker shades = highest. Orthologous pairs have been indicated.

**S1 Video. Time-lapse video showing cytopathic effect in rLA3490-treated HeLa cells as early as 50 min post-exposure**. Time-lapse images (40 frames, 5 s intervals) of rLA3490-treated HeLa cells, 0 to 4 h post-exposure showing noticeable cytopathic effect as early as 50 minutes post-exposure. rLA3490 was added at a final concentration of 5 *µ*g/mL. Images were captured using a Leica DMi8 inverted microscope at 40x magnification. Scale bar 10 *µ*m.

**S2 Video. Time-lapse video of untreated HeLa cells up to four hours post-exposure**. Time-lapse images (40 frames, 5 s intervals) of untreated HeLa cells (control) 0 to 4 h post-exposure showing no morphological changes. Images were captured using a Leica DMi8 inverted microscope at 40x magnification. Scale bar 10 *µ*m.

**S3 Video. Orthogonal projections of t3490-treated HeLa cells, 30 min post-exposure**. Top left and bottom right panels show binding of mCherry-t3490 fusion protein (red) at HeLa cell surface 30 minutes post-exposure. Plasma membrane and nucleus stained with CellMask™ Green Plasma Membrane Stain and ProLong™ Gold Antifade Mountant with DAPI (blue), respectively. Images captured using a Leica SP8 Gated STED 3X super-resolution confocal microscope. Magnification 100x.

**S4 Video. Orthogonal projections of t3490-treated HeLa cells, 60 min post-exposure**. Top left and bottom right panels show binding of mCherry-t3490 fusion at HeLa cell surface 60 minutes post-exposure. Plasma membrane and nucleus stained with CellMask™ Green Plasma Membrane Stain and ProLong™ Gold Antifade Mountant with DAPI, respectively. Images captured using a Leica SP8 Gated STED 3X super-resolution confocal microscope. Magnification 100x.

**S5 Video. Z-stacks of t3490-treated HeLa cells, 30 min post-exposure**. Animation of 24 cross-sectional images (total depth, 6.87 *μ*m; separation, 298.5 nm), with binding of mCherry-t3490 fusion evident at HeLa cell surface. Cells were washed twice with PBS, and the plasma membrane and nucleus stained with CellMask™ Green Plasma Membrane Stain and ProLong™ Gold Antifade Mountant with DAPI, respectively. Images captured using a Leica SP8 Gated STED 3X super-resolution confocal microscope. Magnification 100x.

**S6 Video. Z-stacks of t3490-treated HeLa cells, 60 min post-exposure**. Animation of 32 cross-sectional images (total depth, 9.25 *μ*m; separation, 298.5 nm), with binding of mCherry-t3490 fusion still evident on HeLa cell surface (i.e. no internalization). Cells were washed twice with PBS, and plasma membrane and nucleus stained with CellMask™ Green Plasma Membrane Stain and ProLong™ Gold Antifade Mountant with DAPI, respectively. Images captured using a Leica SP8 Gated STED 3X super-resolution confocal microscope. Magnification 100x.

**S7 Video. Orthogonal projections of HeLa cells treated with rLA3490, 30 min post-exposure**. mCherryr-LA3490 fusion on the HeLa cell surface 30 minutes of post-exposure, with internalization evident (top left, runtime 3 s; and bottom right, runtime 10 s). Plasma membrane and nucleus were stained with CellMask™ Green Plasma Membrane Stain and ProLong™ Gold Antifade Mountant with DAPI, respectively. Images were captured using a Leica SP8 Gated STED 3X super-resolution confocal microscope. Magnification 100x.

**S8 Video. Orthogonal projections of HeLa cells treated with rLA3490, 60 min post-exposure**. Internalized mCherryr-LA3490 fusion evident in all four images. Plasma membrane and nucleus were stained with CellMask™ Green Plasma Membrane Stain and ProLong™ Gold Antifade Mountant with DAPI, respectively. Images were captured using a Leica SP8 Gated STED 3X super-resolution confocal microscope. Magnification 100x.

**S9 Video. Z-stacks of HeLa cells treated with rLA3490, 30 min post-exposure**. Animation of 35 cross-sectional images (total depth, 10.15 *μ*m; separation, 298.5 nm) showing binding of mCherry-rLA3490 fusion to HeLa cell surface. Plasma membrane and nucleus were stained with CellMask™ Green Plasma Membrane Stain and ProLong™ Gold Antifade Mountant with DAPI, respectively. Images were captured using a Leica SP8 Gated STED 3X super-resolution confocal microscope. Magnification 100x.

**S10 Video. Z-stacks of HeLa cells treated with rLA3490, 60 min post-exposure**. Animation of 42 cross-sectional images (total depth, 12.24 *μ*m; separation, 298.5 nm) showing cell membrane binding and internalization of mCherry-rLA3490 fusion. Plasma membrane and nucleus stained with CellMask™ Green Plasma Membrane Stain and ProLong™ Gold Antifade Mountant with DAPI, respectively. Images captured using a Leica SP8 Gated STED 3X super-resolution confocal microscope. Magnification 100x.

